# LncRNA GAS6-AS1 contributes to 5-fluorouracil resistance in colorectal cancer

**DOI:** 10.1101/2024.01.30.577984

**Authors:** Zhonglin Zhu, Minghan Li, Junyong Weng, Shanbao Li, Tianan Guo, Yang Guo, Ye Xu

## Abstract

5-Fluorouracil (5-FU) resistance has always been a formidable obstacle in the adjuvant treatment of advanced colorectal cancer (CRC). In recent years, long non-coding RNAs have emerged as key regulators in various pathophysiological processes including 5-FU resistance. Here, RNA-seq combined with weighted gene correlation network analysis confirmed the close association of GAS6-AS1 with TRG grades. GAS6-AS1 expression was positively correlated with advanced clinicopathological features and poor prognosis in CRC. GAS6-AS1 increased the 50% inhibiting concentration of 5-FU, enhanced cell proliferation, and accelerated G1/S transition in CRC cells, both with and without 5-FU, both in vitro and in vivo. Mechanistically, GAS6-AS1 enhanced the stability of MCM3 mRNA by recruiting PCBP1, consequently increasing MCM3 expression. Furthermore, PCBP1 and MCM3 counteracted the effects of GAS6-AS1 on 5-FU resistance. Notably, the PDX model indicated that combining chemotherapeutic drugs with GAS6-AS1 knockdown yielded superior outcomes in vivo. Together, our findings elucidate that GAS6-AS1 directly binds to PCBP1, enhancing MCM3 expression and thereby promoting 5-FU resistance. GAS6-AS1 may serve as a robust biomarker and potential therapeutic target for combination therapy in CRC.

## Introduction

Colorectal cancer (CRC) ranks as the third most prevalent cancer globally in terms of incidence and is the second leading cause of cancer-related mortality(Siegel, Miller et al., 2023, Sung, Ferlay et al., 2021). Owing to the subtle symptoms associated with CRC, a majority of patients receive a diagnosis at an advanced stage, resulting in a 5-year survival rate of less than 20% for metastatic CRC(Siegel, Miller et al., 2020). In recent years, there has been a continuous expansion of research into the comprehensive treatment of CRC(Weng, Li et al., 2022). Adjuvant chemotherapy has evolved as an integral component of the comprehensive CRC treatment approach, wherein the formidable challenge of chemotherapy resistance has become apparent(Allen & Johnston, 2005, Wilson, Danenberg et al., 2014). The 5-fluorouracil (5-FU)-based chemotherapy regimen serves as the primary therapeutic treatment for adjuvant treatment in CRC(Vodenkova, Buchler et al., 2020). However, the precise molecular mechanisms underlying 5-FU resistance remain inadequately elucidated.

Long non-coding RNAs (lncRNAs) encompass a class of RNA molecules exceeding 200 nucleotides in length, distinguishing them from small non-coding RNAs(Chen & Shen, 2020, Chen, Lin et al., 2020, Liu, Dang et al., 2021). Devoid of protein-coding functions, lncRNAs actively participate in gene regulation in the RNA form(Adnane, Marino et al., 2022, Nemeth, Bayraktar et al., 2023). Substantial research has revealed that lncRNAs can modulate gene expression and molecular pathways through various mechanisms, primarily serving as signals, decoys, guides, and scaffolds(Statello, Guo et al., 2021, Wang & Chang, 2011). Significantly, an expanding body of evidence suggests that lncRNAs play an indispensable role in the progression of various cancer types, influencing aspects such as proliferation, metastasis, chemoradiotherapy resistance and immunosuppression. LINC01138 is highly expressed in hepatocellular carcinoma tissues and promotes cell proliferation and metastasis(Li, Zhang et al., 2018). LncRNA XIST boosts proliferation and self-renewal of cancer stem cells in breast cancer through IL-6/STAT3 signaling(Ma, Zhu et al., 2023). Additionally, lncRNA LUCAT1 is overexpressed in CRC tissues and contributes to cell growth, as well as resistance to 5-FU and Oxaliplatin(Huan, Guo et al., 2020). A recently discovered lncRNA, DILA1, has been found to stimulate cell growth, facilitate cell cycle G1/S progression and induce tamoxifen resistance in breast cancer by inhibiting Cyclin D1 degradation(Shi, Li et al., 2020). Nevertheless, the intricate roles and molecular mechanisms of lncRNAs in the context of 5-FU resistance remain elusive(Liu, Gao et al., 2020, Wei, Sun et al., 2020).

In this study, the integration of RNA-seq with weighted gene correlation network analysis (WGCNA) validated the close correlation of a specific lncRNA with tumor regression grading (TRG) in CRC. TRG, in accordance with the guidelines of the American Joint Committee on Cancer (AJCC), serves as a pivotal pathological evaluation indicator for tumors post-neoadjuvant chemotherapy. It facilitates the quantitative analysis of the therapeutic efficacy of chemotherapy drugs on tumors. Our findings revealed that GAS6-AS1 not only heightened cell growth, G1/S transition and the 50% inhibiting concentration (IC50) of 5-FU, but also functioned mechanistically as a ‘guider.’ GAS6-AS1 increased the binding of PCBP1 with MCM3 mRNA, consequently promoting the expression of MCM3. Notably, clinical correlation analysis and insights from the patient-derived xenograft (PDX) model collectively suggested that GAS6-AS1 could serve as a robust biomarker and a promising therapeutic target for combination drug therapy in CRC.

## Results

### GAS6-AS1 is closely associated with 5-FU resistance and might serve as a sensitive indicator for predicting patient prognosis in CRC

To investigate the key regulators in chemotherapy resistance of CRC, RNA-seq analysis (TRG 0-1 *vs* TRG 3) was performed to reveal the differentially expressed lncRNAs (Fig 1a) and mRNAs (Fig 1b). Then, we performed WGCNA to explore key regulators closely associated with TRG stage (Fig 1c and EV1a). 18 modules were identified to be correlated with TRG trait, of which the MEblue module is the most closely associated module (Fig 1d and EV1b). To screen the hub genes in the MEblue module, we drew the profile of module membership vs gene significance, and highlighted the most significant part of hub genes (Fig 1e). Then, we constructed comprehensive network analysis with these hub genes (Fig 1f). We focused on the four lncRNAs (Fig 1g), and found that both GAS6-AS1 and CATIP-AS1 were highly expressed in CRC tissues from TCGA database (Fig 1h, EV1c and e). In addition, patients with high GAS6-AS1 showed poor overall survival though there was no significant statistical difference (EV1d), while CATIP-AS1 showed opposite effect (EV1f). We didn’t find another two lncRNAs in TCGA database. Thereby, we focused on GAS6-AS1 in following study. Then, we retrieved the CRC data from GEO database. GAS6-AS1 was highly expressed in CRC tissues (GSE39582) and positively correlated with poor disease-free survival (GSE106584) (EV1g and h).

**Figure 1.**
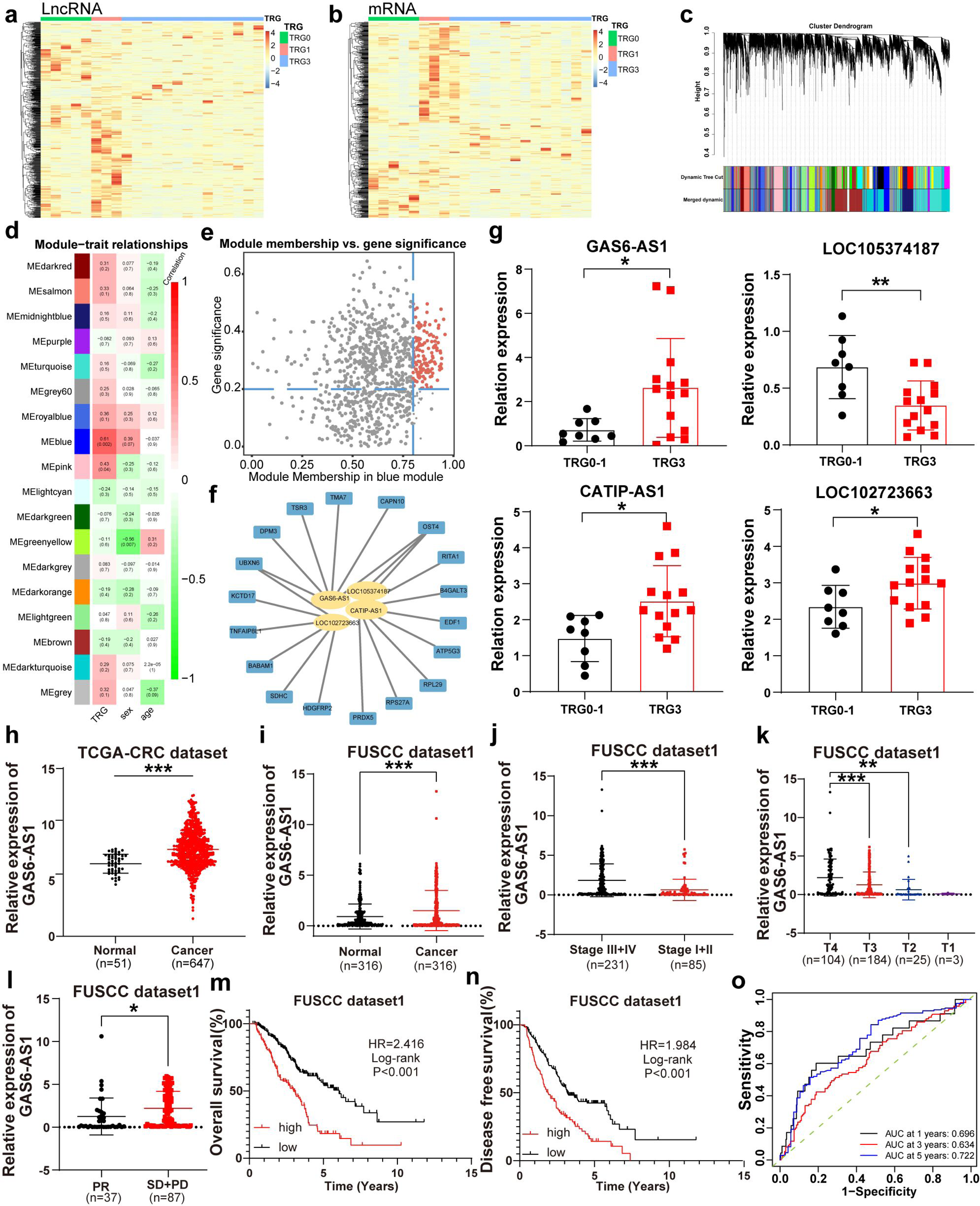
GAS6-AS1 is closely associated with 5-FU resistance and might serve as a sensitive indicator for predicting patient prognosis in CRC. **a** The lncRNA heatmap of RNA-seq between TRG 0-3. **b** The mRNA heatmap of RNA-seq between TRG 0-3. **c** The gene cluster dendrogram and dynamic module color of WGCNA. **d** The relationship between modules and traits. **e** The profile of module membership vs gene significance. **f** The comprehensive network of hub genes. **g** The expression of GAS6-AS1 (Fold change=3.65, P=0.0293), LOC105374187 (Fold change=0.51, P=0.0293), CATIP-AS1 (Fold change=1.28, P=0.0408) and LOC102723663 (Fold change=1.70, P=0.0150) between samples with TRG 0-1 and TRG3. **h** The level of GAS6-AS1 between CRC samples (n=647) of TCGA database compared with corresponding normal tissues (n=51). **i** The level of GAS6-AS1 in CRC samples and paired adjacent normal tissues (n=316) of FUCSC dataset. **j** The level of GAS6-AS1 is higher in CRC samples with stage III and IV (n=231) than these with stage I and II (n=85). **k** The level of GAS6-AS1 gradually decreases in the samples with the stage from T4 to T1. **l** The level of GAS6-AS1 is lower in the samples with partial response (PR) of distant metastatic lesion than these with stable disease (SD) and progressive disease (PD) of distant metastatic lesion. **m** Kaplan-Meier analysis shows that the high level of GAS6-AS1 correlates with poorer overall survival in all patients. **n** The patients with high level of GAS6-AS1 show a poor disease free survival in all patients. **o** ROC analysis confirms that GAS6-AS1 level is a sensitive indicator for predicting patient prognosis. *P < 0.05; **P < 0.01; ***P < 0.001.

GAS6-AS1 is located in 13q34 with the length of 902 bp (EV2a). Along with the thorough research, it has been discovered that a small number of lncRNAs can be translated into peptides. Therefore, we next analyzed the protein-coding potential of GAS6-AS1with four independent mathematical methods, including Coding Potential Assessment Tool(Wang, Park et al., 2013), Coding Potential Calculator(Kong, Zhang et al., 2007), length and guanine-cytosine (LGC) algorithms(Wang, Yin et al., 2019) and ORF finder software from NCBI (EV2b-e). All these methods showed that GAS6-AS1 didn’t have protein-coding potential. The Ensemble database indicated that GAS6-AS1 shared no homology with other genomic regions. The secondary structure of GAS6-AS1 was listed in Supplementary EV2f.

Next, the clinical significance of GAS6-AS1 were analyzed. GAS6-AS1 level was up-regulated in 55.38% (175/316) CRC tissues (Fig 1i), and positively correlated with differentiation grade, T stage, N stage, M stage and TNM stage (Fig 1j and k, EV3a-g, Table EV1). To analyze the correlations of GAS6-AS1 expression with objective response rate (ORR) of 5-FU based chemotherapy regimen, the chemotherapy response was evaluated in 138 cases with distant metastasis of 316 patients. The ORR reached 29.84% and the level of GAS6-AS1 was higher in cases with stable disease (SD) and progressive disease (PD) than these with partial response (PR) (Fig 1l). Kaplan-Meier analysis showed that the high level of GAS6-AS1 correlated with poorer overall survival and disease-free survival in all patients (Fig 1m-n). Then, patients were stratified based on clinicalpathological characteristics. Patients with high level of GAS6-AS1 in stage III+IV, T4, T1-3, M1 and M0 all showed poorer overall survival and disease-free survival (Fig EV3h-m). More importantly, the Hazard Ratio (HR) value in patients with stage III+IV, T4 and M1 were higher than these with stage I+II, T1-3 and M0, respectively. Univariate and multivariate regression analysis indicated that GAS6-AS1 level is an independent prognostic factor for CRC (Table EV2). Also, receiver operating characteristic curve (ROC) analysis confirmed that GAS6-AS1 level was a sensitive indicator for predicting patient prognosis, especially for 5 years overall survival (Fig 1o).

### GAS6-AS1 promotes 5-FU resistance and malignant behaviors in CRC cells in vitro

To further elucidate the roles of GAS6-AS1 in 5-FU resistance, we firstly constructed CRC cell lines with 5-FU resistance. The results showed that IC50 value was significantly higher in HCT15-Re (Fold change=5.26) and HCT8-Re (Fold change=3.74) cells compared with HCT15 and HCT8 cells (Fig 2a-b). Secondly, RT-PCR results indicated that the expression of GAS6-AS1 was significantly higher in both HCT15-Re and HCT8-Re cells than that in HCT15 and HCT8 cells (Fig 2c). Thirdly, we generated stable overexpression cell lines of GAS6-AS1 with HCT15 and HCT8 cells, and stable knockdown cell lines of GAS6-AS1 with HCT15-Re and HCT8-Re cells (Fig EV4a and b). Then, the effects of GAS6-AS1 on 5-FU resistance was explored. GAS6-AS1 significantly increased the IC50 of 5-FU, while knockdown of GAS6-AS1 decreased it, in CRC cells (Fig 2d-g). Next, CCK8, colony formation, edu and flow cytometry for cell cycle assays confirmed that overexpression of GAS6-AS1 promoted cell growth and cell cycle G1/S transition with or without 5-FU, while knockdown of GAS6-AS1 exerted the opposite effects, in HCT15 and HCT8 cells (Fig 2h-o and EV4c-j). Collectively, these data implied that GAS6-AS1 might enhanced 5-FU resistance and cell growth in CRC cells through promoting cell cycle G1/S progression.

**Figure 2.**
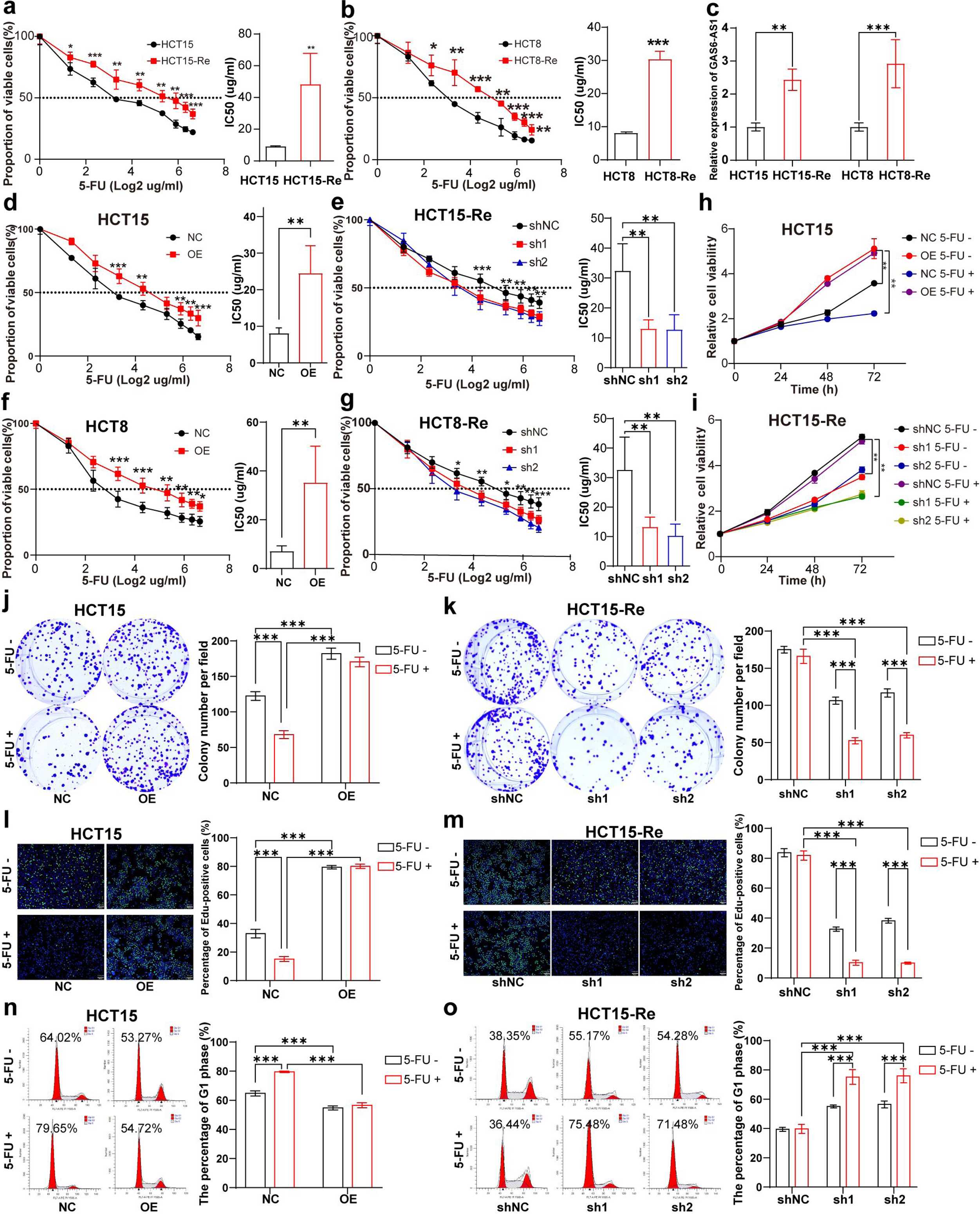
GAS6-AS1 promotes 5-FU resistance and malignant behaviors in CRC cells in vitro. **a** The IC50 value between HCT15 and HCT15-Re cells. **b** The IC50 value between HCT8 and HCT8-Re cells. **c** The expression of GAS6-AS1 between HCT15 and HCT15-Re cells, HCT8 and HCT8-Re cells. **d** The IC50 value in HCT15 with NC or overexpression of GAS6-AS1. **e** The IC50 value in HCT15-Re cells with shNC or shRNA target GAS6-AS1. **f** The IC50 value in HCT8 with NC or overexpression of GAS6-AS1. **g** The IC50 value in HCT8-Re cells with shNC or shRNA target GAS6-AS1. **h-i** The CCK8 results in HCT15 and HCT15-Re cells with or without 5-FU. **j-k** The colony formation results in HCT15 and HCT15-Re cells with or without 5-FU. **l-m** The edu results in HCT15 and HCT15-Re cells with or without 5-FU. **n-o** The flow cytometry for cell cycle assays in HCT15 and HCT15-Re cells with or without 5-FU. *P < 0.05; **P < 0.01; ***P < 0.001.

### GAS6-AS1 promotes 5-FU resistance in vivo

Furthermore, the effects of GAS6-AS1 on 5-FU resistance were observed in xenograft mouse models. Overexpression of GAS6-AS1 increased the volume and weight of tumors under the treatment of 5-FU, while knockdown of GAS6-AS1 decreased those under the treatment of 5-FU, in HCT15 and HCT8 cells (Fig 3a-l). Then, we detected the expression of MCM3, KI67 (biomarkers of cell proliferation), cyclin D1 and CDK4 (biomarkers of cell cycle) in continuous tumor tissue slice. Overexpression of GAS6-AS1 increased the expression of these genes under the treatment of 5-FU, while knockdown of GAS6-AS1 decreased those under the treatment of 5-FU, in HCT15 and HCT8 cells (Fig 3m). Consequently, GAS6-AS1 promotes 5-FU resistance of CRC cells in vivo.

**Figure 3.**
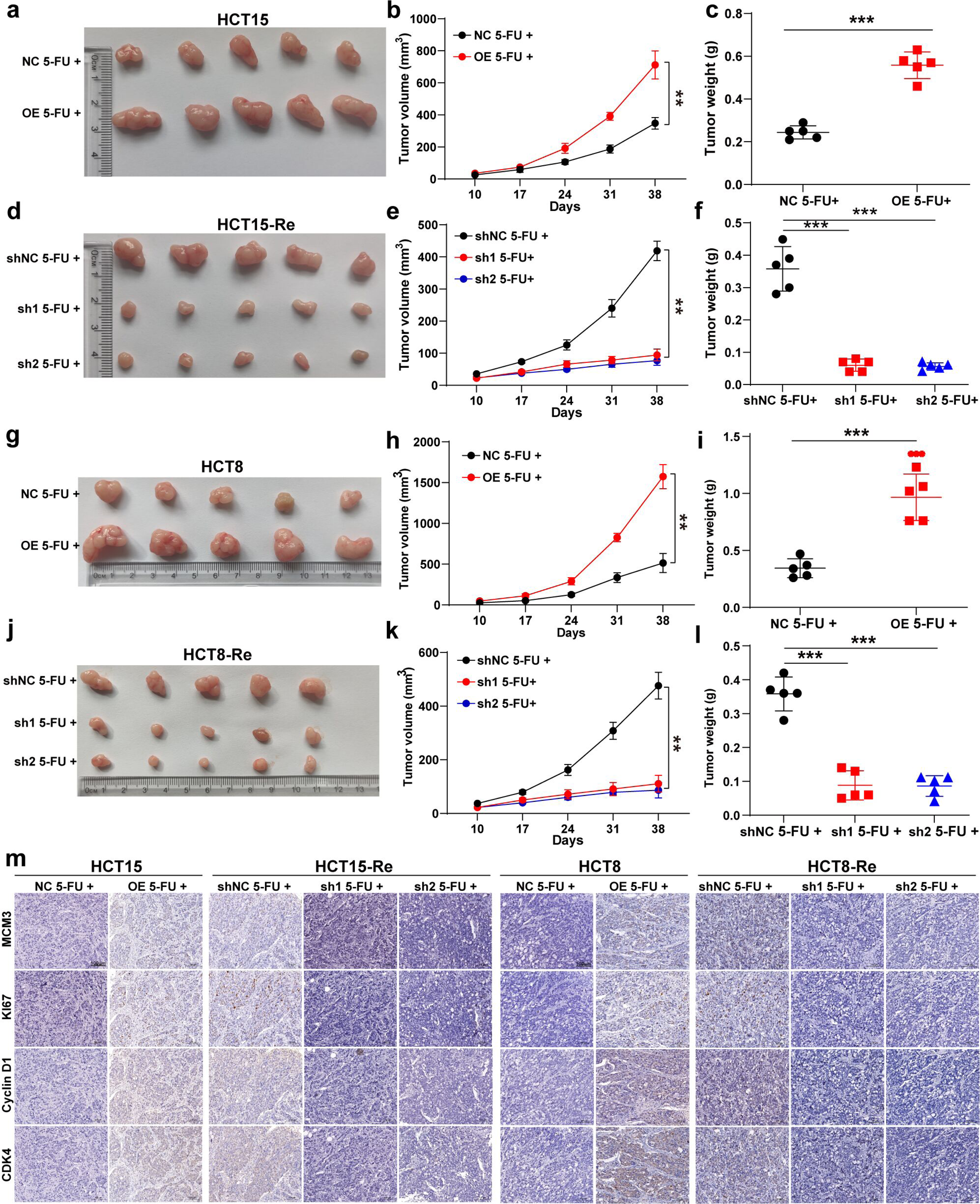
GAS6-AS1 promotes 5-FU resistance in vivo. **a** Images of subcutaneous xenograft tumors of HCT15 cells. **b** Tumor volumes of HCT15 cells were measured every 7 days. **c** Tumor weights of HCT15 cells were shown. **d** Images of subcutaneous xenograft tumors of HCT15-Re cells. **e** Tumor volumes of HCT15-Re cells were measured every 7 days. **f** Tumor weights of HCT15-Re cells were shown. **g** Images of subcutaneous xenograft tumors of HCT15 cells. **h** Tumor volumes of HCT15 cells were measured every 7 days. **i** Tumor weights of HCT15 cells were shown. **j** Images of subcutaneous xenograft tumors of HCT15-Re cells. **k** Tumor volumes of HCT15-Re cells were measured every 7 days. **l** Tumor weights of HCT15-Re cells were shown. **m** Representative images of IHC results of MCM3, KI67, Cyclin D1 and CDK4 in each group. *P < 0.05; **P < 0.01; ***P < 0.001.

### GAS6-AS1 physically interacts with PCBP1 in CRC cells

Studies have revealed that the molecular mechanism of lncRNAs diversifies according to its subcellular distribution. Both FISH and nuclear cytoplasmic separation assays indicated that most of GAS6-AS1 located in cytoplasm (Fig 4a-b). In addition, most of the roles of lncRNAs require interactions with one or more RNA-binding proteins (RBPs). Next, we dedicated to seeking for the RBPs of GAS6-AS1. RNA pulldown and mass spectrometry were performed in HCT15 and HCT8 cells (Fig 4c). With the criterions of molecular weight between 20-55kDa, more than 3 peptides and more than 3 unique peptides, five proteins existed between the intersections of the results of HCT15 and HCT8 (Fig 4d). After checking the features of those proteins, hnRNPK and PCBP1 were selected for further research. Only PCBP1 protein was pulled down by GAS6-AS1 sense probe (Fig 4e). Moreover, RIP assays confirmed the association between GAS6-AS1 with PCBP1 (Fig 4f). As expected, there was no enrichment of MALAT1 in each group. According to the binding motif of PCBP1 (Fig 4g), three potential binding sites in GAS6-AS1 was found. Threes truncated RNA separately containing three binding sites and full-length RNA with biotin labeling were constructed according to the secondary structure of GAS6-AS1 (Fig 4h and i). RNA pulldown assays confirmed that the binding site 2 in GAS6-AS1 was required for the interaction with PCBP1, not the binding site 1 and 3 (Fig 4j). Furthermore, three Flag-tagged truncated PCBP1 and full length were generated based on the functional domains (Fig 4k). RIP assays showed that deletion mutation of KH3 abolished its interaction with GAS6-AS1 (Fig 4l). In a word, we demonstrated that GAS6-AS1 physically interacted with PCBP1 in CRC cells through binding site 2 and KH3.

**Figure 4.**
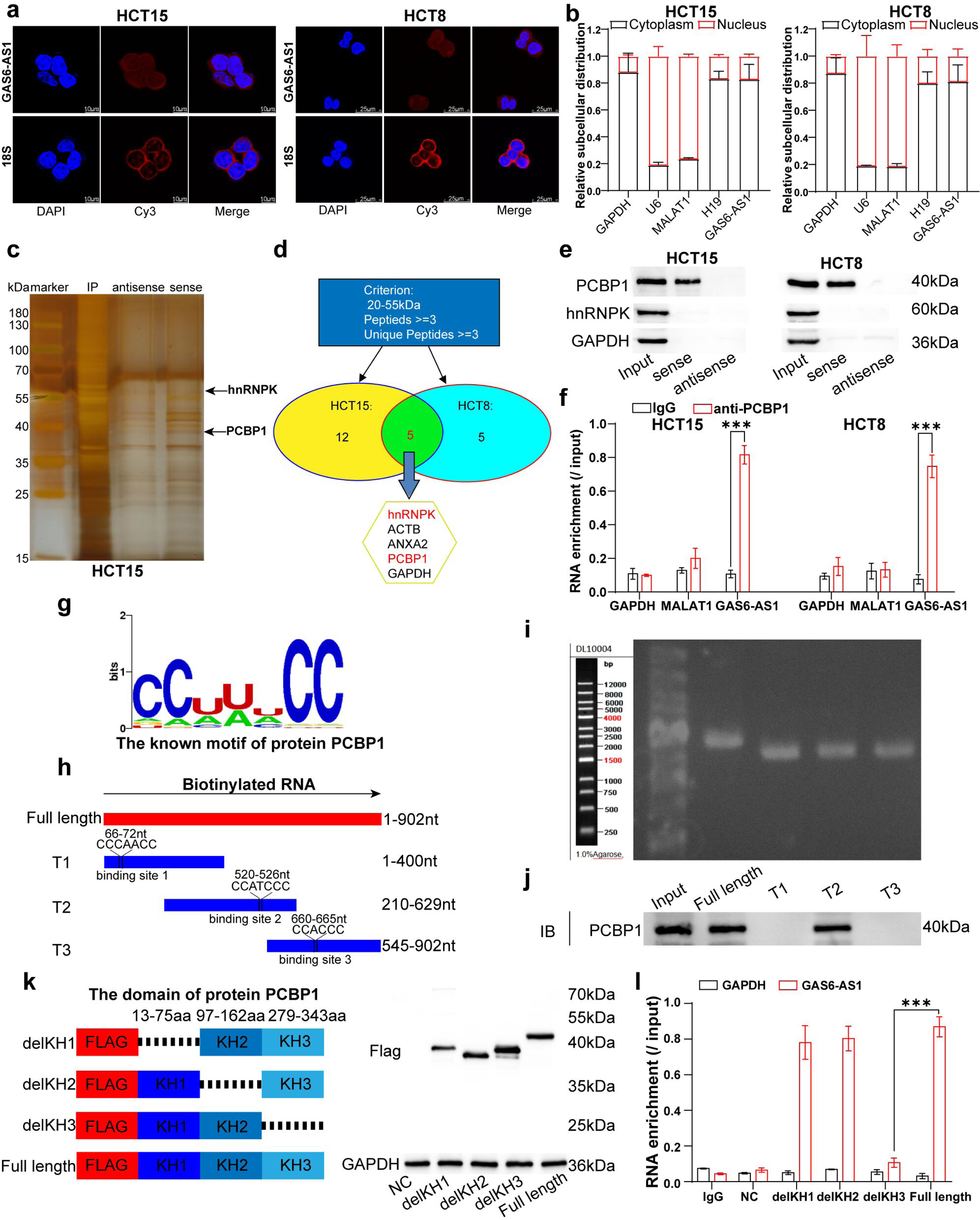
GAS6-AS1 physically interacts with PCBP1 in CRC cells. **a** The representative images of RNA-FISH with GAS6-AS1 probe in HCT15 and HCT8 cells. **b** The results of nuclear cytoplasmic separation assays in HCT15 and HCT8 cells. **c** The silver staining of proteins pulled down by GAS6-AS1 probe in HCT15 cells. **d** The filter criterion of mass spectrometry results. **e** The RNA pulldown results of GAS6-AS1 sense and antisense probe in HCT15 and HCT8 cells. **f** The RIP assays with anti-PCBP1 in HCT15 and HCT8 cells. **g** The binding motif of PCBP1. **h** The design drawing of truncated RNA. **i** The gel electrophoresis map of truncated RNA. **j** The RNA pulldown results with truncated RNA of GAS6-AS1 in HCT15 cells. **k** The design drawing and western blot testing of truncated PCBP1 protein. **l** The RIP assays with anti-Flag in HCT15 cells transfected with the above truncated PCBP1 plasmids. ***P < 0.001.

### PCBP1 is required for GAS6-AS1 in the role of 5-FU resistance

In order to explore the role of PCBP1 in the regulation of GAS6-AS1 on 5-FU resistance, we generated overexpression plasmid and threes shRNAs targeted PCBP1 (Fig 5a). CCK8 results showed that knockdown of PCBP1 rescued the enhance of GAS6-AS1 on the IC50 of 5-FU, while overexpression of PCBP1 increased the IC50 of 5-FU decreased by downregulation of GAS6-AS1 (Fig 5b and c). Furthermore, CCK8, colony formation, edu and flow cytometry for cell cycle assays elucidated that PCBP1 could rescue the regulation of GAS6-AS1 on the cell growth and cell cycle G1/S transition under the treatment of 5-FU (Fig 5d-g). In brief, PCBP1 is required for GAS6-AS1 in the promotion of 5-FU resistance, cell growth and cell cycle G1/S progression in CRC.

**Figure 5.**
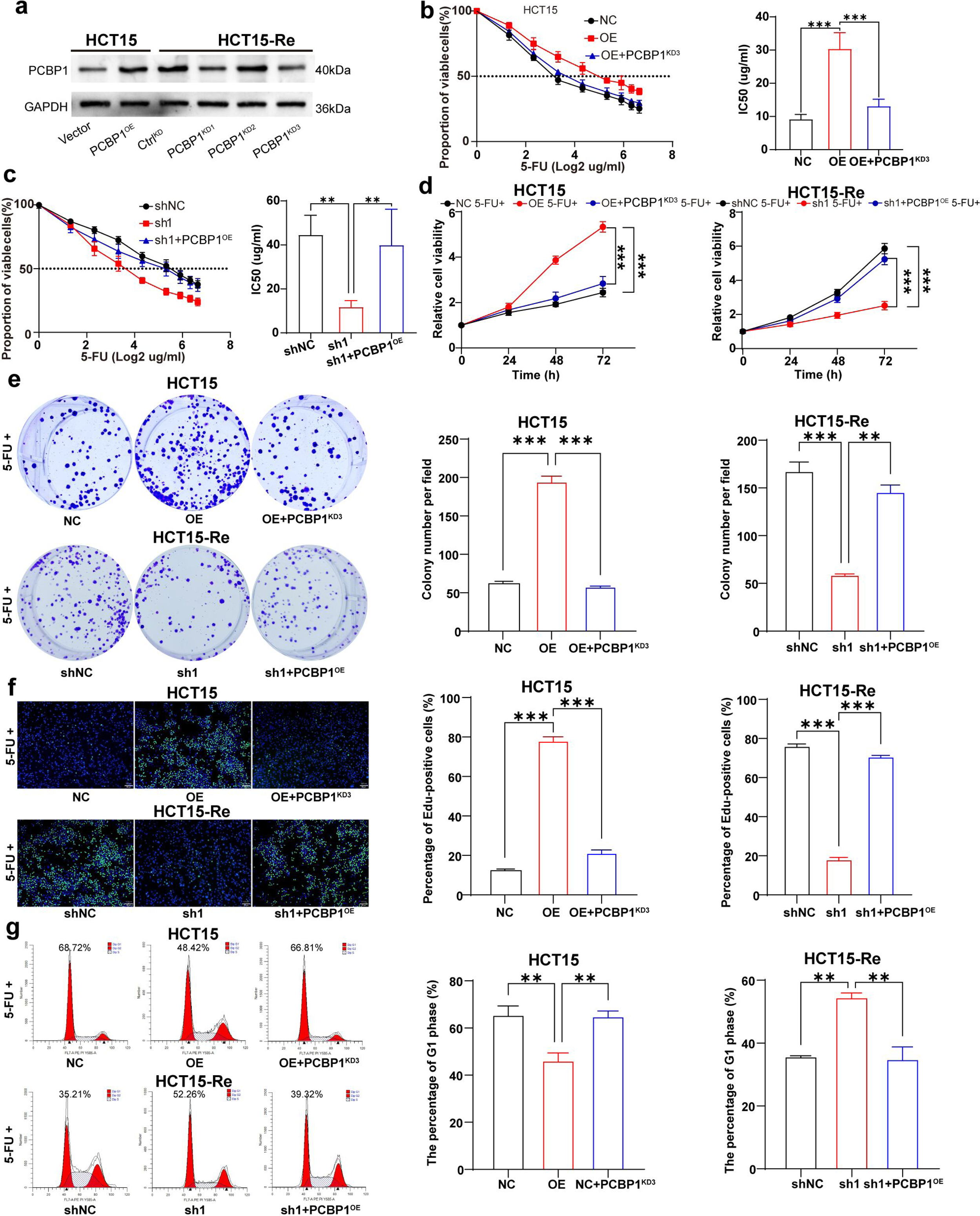
PCBP1 is required for GAS6-AS1 in the role of 5-FU resistance. **a** Western blot tested the efficacy of overexpression plasmid and threes shRNAs targeted PCBP1. **b-c** The IC50 value in HCT15 and HCT15-Re cells with alteration of GAS6-AS1 and PCBP1. **d** The CCK8 results in HCT15 and HCT15-Re cells with alteration of GAS6-AS1 and PCBP1 under the treatment of 5-FU. **e** The colony formation results in HCT15 and HCT15-Re cells with alteration of GAS6-AS1 and PCBP1 under the treatment of 5-FU. **f** The edu results in HCT15 and HCT15-Re cells with alteration of GAS6-AS1 and PCBP1 under the treatment of 5-FU. **g** The flow cytometry for cell cycle results in HCT15 and HCT15-Re cells with alteration of GAS6-AS1 and PCBP1 under the treatment of 5-FU. **P < 0.01; ***P < 0.001.

### MCM3 is a functional downstream mediator for GAS6-AS1

To search for the functional downstream of GAS6-AS1, RNA-seq on HCT15 cells with overexpression of GAS6-AS1 was performed (Fig 6a). In addition, two GEO datasets were selected: GSE16236 (RNA-seq on HCT116 cells with knockdown of PCBP1) and GSE131210 (CLIP-seq on HCT116 cells with anti-PCBP1) (Fig 6a). There were 17 genes in the intersect of the three datasets (Fig 6b). GO analysis indicated that those genes mainly enriched in the biological processes related with DNA replication (Fig 6c). Furthermore, the 22 RNA sequencing samples mentioned above were classified into high GAS6-AS1group and low GAS6-AS1 group according to the mean of GAS6-AS1 level. GSEA analysis showed that high GAS6-AS1 participated in DNA replication, DNA repair, DNA damage repair signal transduction, DNA biosynthetic process and DNA binding (Fig EV5). MCM3, MCM4, PARD6B, BRIP1 and RAD54B were enriched in the above biological process. RT-PCR results elucidated that only MCM3 mRNA level upregulated after overexpression of GAS6-AS1 or PCBP1, and downregulated after knockdown of GAS6-AS1 or PCBP1 (Fig 6d and f). West blot also confirmed the results (Fig 6e and g). Meanwhile, both GAS6-AS1 and PCBP1 promoted the expression of cyclin D1 and CDK4 (Fig 6e and g).

**Figure 6.**
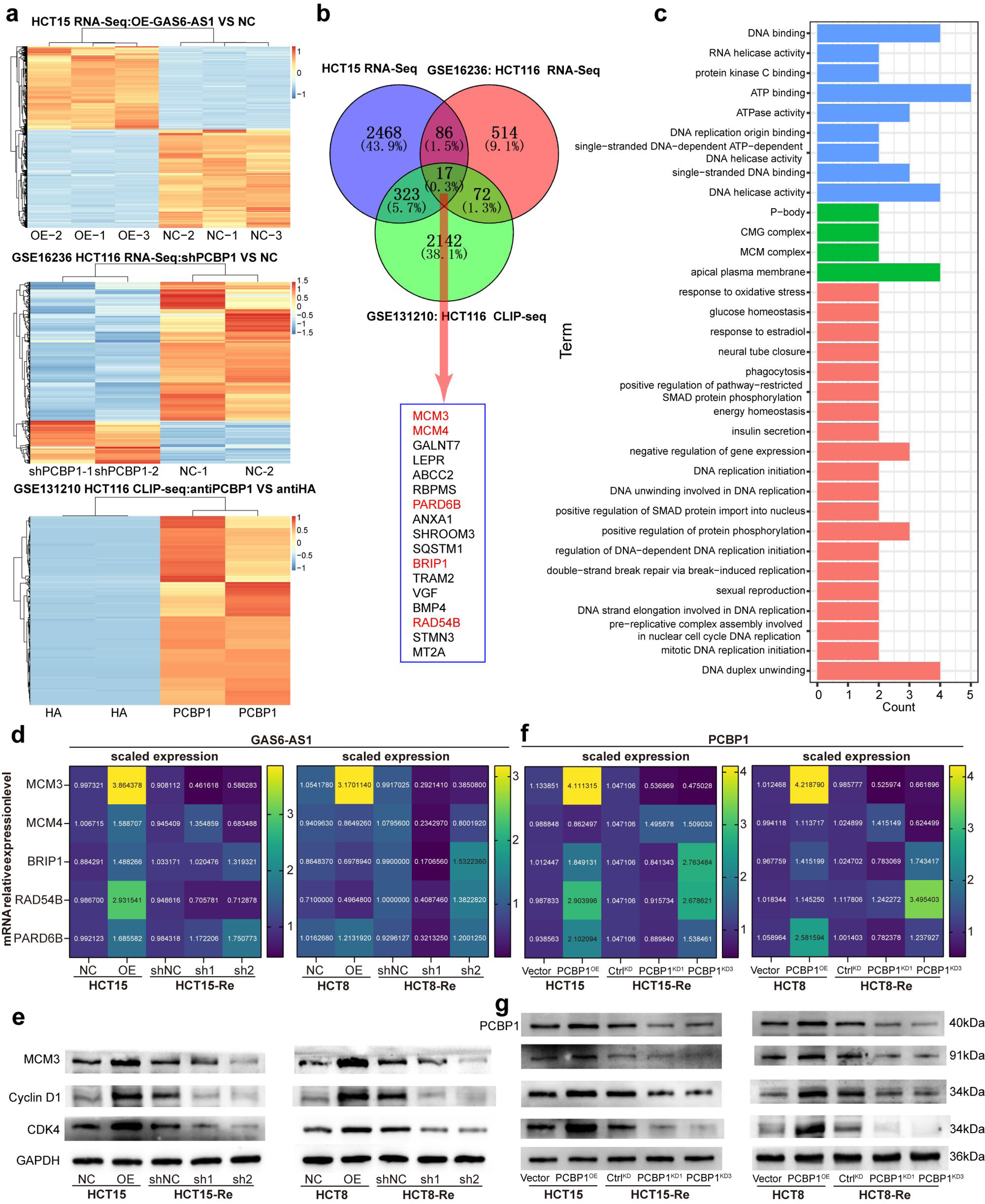
MCM3 is a functional downstream mediator for GAS6-AS1. **a** The heatmap of RNA-seq on HCT15 cells with overexpression of GAS6-AS1, the RNA-seq on HCT116 cells with knockdown of PCBP1 in GSE16236 dataset and the CLIP-seq on HCT116 cells with anti-PCBP1 in GSE131210 dataset. **b** The intersect genes in the above three RNA-seq data. **c** The GO analysis of the above intersect genes. **d** The RT-PCR results of five genes in HCT15 and HCT8 cells with overexpression or knockdown of GAS6-AS1. **e** The western blot results of five genes in HCT15 and HCT8 cells with overexpression or knockdown of GAS6-AS1. **f** The RT-PCR results of five genes in HCT15 and HCT8 cells with overexpression or knockdown of PCBP1. **g** The western blot results of five genes in HCT15 and HCT8 cells with overexpression or knockdown of PCBP1.

To explore whether MCM3 could act as the downstream mediator of GAS6-AS1, rescue experiments were employed. Overexpression plasmid and threes shRNAs targeted MCM3 were constructed (Fig EV6a). CCK8, colony formation, edu and flow cytometry for cell cycle assays confirmed that MCM3 could rescue the regulation of GAS6-AS1 on the 5-FU resistance, cell growth and cell cycle G1/S progression (Fig EV6b-g). In summary, MCM3 is required for GAS6-AS1 in the role of 5-FU resistance, cell growth and cell cycle G1/S progression in CRC.

### GAS6-AS1 enhances the stability of MCM3 mRNA through increasing its binding to PCBP1

To analyze the interaction between PCBP1 and MCM3, RNA pulldown and RIP assay were performed. PCBP1 protein was pulled down by MCM3 mRNA sense probe but not antisense (Fig 7a). Moreover, RIP assays confirmed the binding between MCM3 mRNA with PCBP1 (Fig 7b and c). According to the binding motif of PCBP1, three potential binding sites in the 3’-UTR of MCM3 mRNA was marked. Threes truncated RNA and full-length RNA with biotin labeling were constructed (Fig 7d). RNA pulldown assays confirmed that the binding site 3 in the 3’-UTR of MCM3 mRNA was required for the interaction with PCBP1, not the binding site 1 and 2 (Fig 7e). Furthermore, RIP assays with the above three Flag-tagged truncated PCBP1 and full length indicated that deletion mutation of KH1 failed to bind to the 3’-UTR of MCM3 mRNA (Fig 7f).

**Figure 7.**
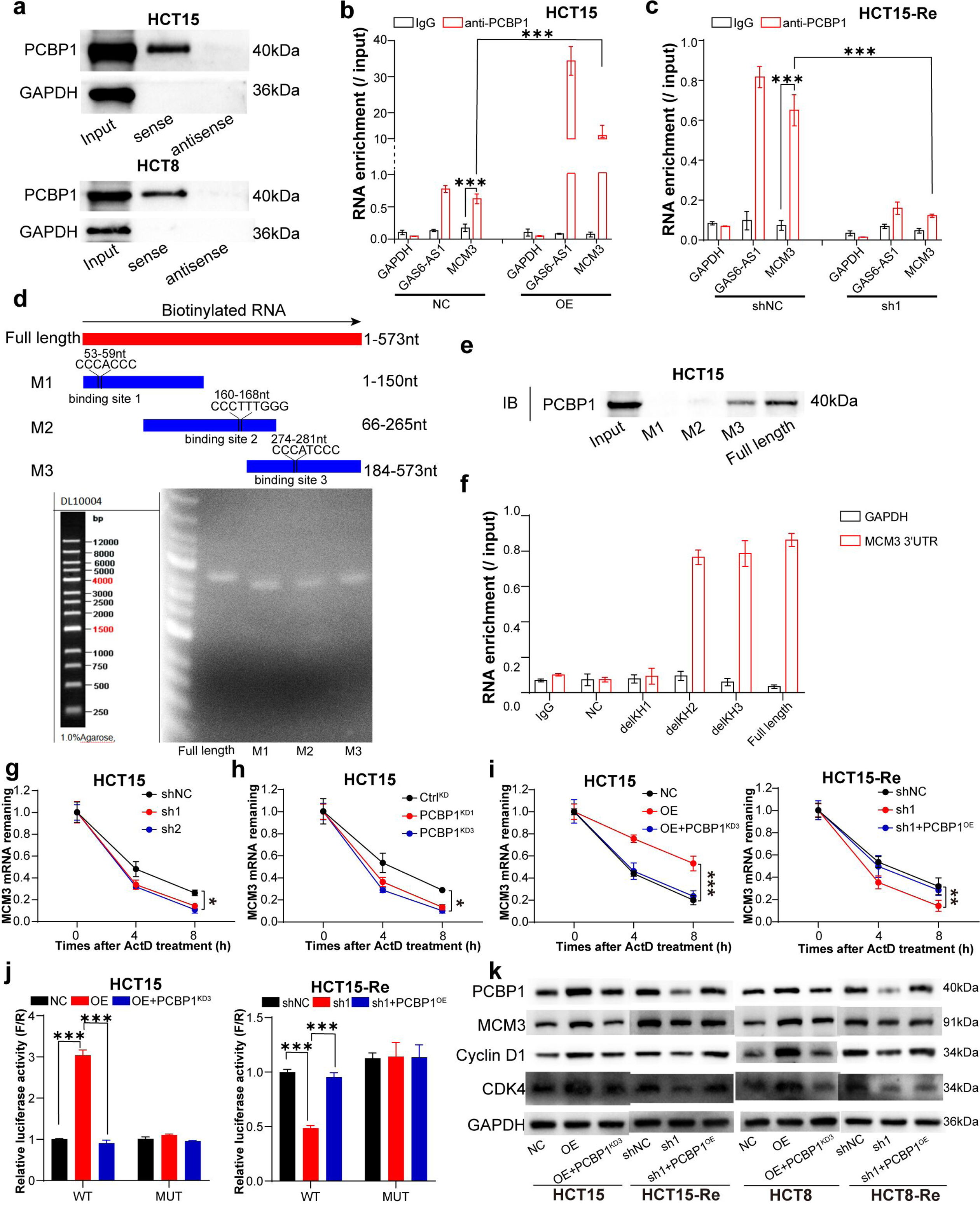
GAS6-AS1 enhances the stability of MCM3 mRNA through increasing its binding to PCBP1. **a** The RNA pulldown assays with MCM3 mRNA probe in HCT15 and HCT8 cells. **b** The RIP assays with anti-PCBP1 in HCT15 cells with overexpression of GAS6-AS1. **c** The RIP assays with anti-PCBP1 in HCT15-Re cells with knockdown of GAS6-AS1. **d** The design drawing of truncated 3’-UTR of MCM3 mRNA. **e** The RNA pulldown assays with truncated 3’-UTR of MCM3 mRNA probe in HCT15 cells. **f** The RIP assays with anti-Flag in HCT15 cells transfected with the truncated PCBP1 plasmids. **g** The relative MCM3 mRNA remaining in HCT15 cells with knockdown of GAS6-AS1 after treatment with ActD. **h** The relative MCM3 mRNA remaining in HCT15 cells with knockdown of PCBP1 after treatment with ActD. **i** The relative MCM3 mRNA remaining in HCT15 and HCT15-Re cells with alteration of GAS6-AS1 and PCBP1 after treatment with ActD. **j** The luciferase reporter plasmids containing wild type of 3’-UTR of MCM3 mRNA or mutant type in binding stie 3 of 3’-UTR of MCM3 mRNA were constructed. Luciferase reporter assays were performed in HCT15 and HCT15-Re cells with alteration of GAS6-AS1 and PCBP1. **k** The western blot results of PCBP1, MCM3, Cyclin D1and CDK4 in HCT15 and HCT8 cells with alteration of GAS6-AS1 and PCBP1. *P < 0.05; **P < 0.01; ***P < 0.001.

As a member of heterogeneous nuclear ribonucleoproteins (hnRNPs), the main roles of PCBP1 include regulation of transcription, control of mRNA stability, translation regulation and alternative splicing. Given that GAS6-AS1 located in cytoplasm and regulated the expression of mRNA and protein of MCM3, we speculated that PCPB1 might regulated the stability of MCM3 mRNA through binding to the 3’-UTR of mRNA. HCT15 cells were treated with Actinomycin D to block de novo transcription. RT-PCR results revealed that knockdown of GAS6-AS1 or PCBP1 both decreased the stability of MCM3 mRNA (Fig 7g and h). Moreover, PCBP1 rescued the regulation of GAS6-AS1 on the stability of MCM3 mRNA (Fig 7i).

To dissect the role of GAS6-AS1 on the binding of PCBP1 on MCM3, further RIP assays were conducted. The enrichment of MCM3 mRNA by PCBP1 was remarkably increased by overexpression of GAS6-AS1, and decreased by knockdown of GAS6-AS1 (Fig 7b and c), suggesting that GAS6-AS1 promoted the interaction between MCM3 mRNA and PCBP1. Meanwhile, we generated luciferase plasmids containing wild type of 3’-UTR of MCM3 mRNA or mutant type in binding stie 3 of 3’-UTR of MCM3 mRNA. GAS6-AS1 significantly enhanced the luciferase activity of wild type, while knockdown of GAS6-AS1 decreased the luciferase activity of wild type, but not the mutant type (Fig 7j). Interestingly, knockdown of PCBP1 decreased the luciferase activity of wild type enhanced by overexpression of GAS6-AS1, while overexpression of PCBP1 increased it which was reduced by knockdown of GAS6-AS1 (Fig 7j). Furthermore, we confirmed that the promotion of MCM3 expression by GAS6-AS1 is dependent on PCBP1 by western blotting (Fig 7k). We observed that knockdown of PCBP1 decreased MCM3 expression enhanced by overexpression of GAS6-AS1, while overexpression of PCBP1 increased it downregulated by knockdown of GAS6-AS1. To sum up, GAS6-AS1 can serve as a ‘guider’ to promote the interaction of PCBP1 and MCM3 mRNA 3’-UTR, thereby enhancing the stability and expression of MCM3 mRNA.

### GAS6-AS1 might serve as a powerful biomarker and potential therapeutic target for combination drug therapy in CRC

To further dissect the clinical significance of GAS6-AS1 in CRC, we detected its expression in our TMA with RNA-FISH. GAS6-AS1 level was up-regulated in 65.82% (208/316) CRC tissues (Fig 8a and d, EV7a). The level of GAS6-AS1 was positively correlated with differentiation grade, T stage, N stage, M stage and TNM stage (Fig 8b, c, e and f, EV7b-h, Table EV3). Kaplan-Meier analysis showed that patients with high level of GAS6-AS1 showed poorer overall survival and disease-free survival in all patients (Fig 8g and h). Then, patients were stratified based on clinicalpathological characteristics. The results showed that HR in patients of different stages was different (Fig EV7i-p). In addition, univariate and multivariate regression analysis indicated that GAS6-AS1 level is an independent prognostic factor for CRC (Table EV4). Also, ROC analysis confirmed that GAS6-AS1 level was a specific indicator for predicting patient prognosis (Fig 8i).

**Figure 8.**
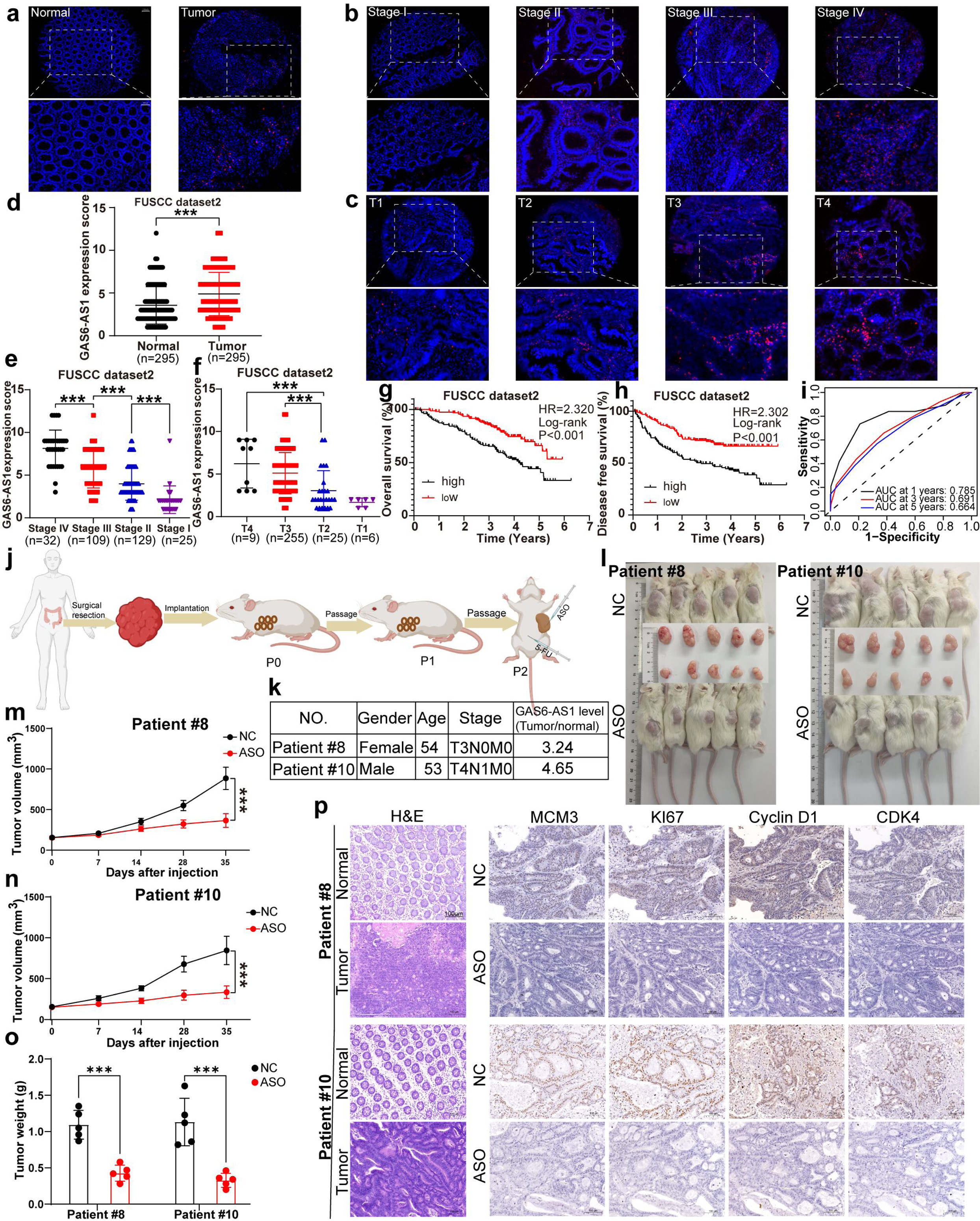
GAS6-AS1 might serve as a powerful biomarker and potential therapeutic target for combination drug therapy in CRC. **a** Representative images of GAS6-AS1 RNA-FISH of normal and tumor tissues in our TMA. **b** Representative images of GAS6-AS1 RNA-FISH of stage I to stage IV in our TMA. **c** Representative images of GAS6-AS1 RNA-FISH of T1 to T4 in our TMA. **d** GAS6-AS1 expression score was significantly higher in CRC tissues than in normal tissues. **e** The expression score of GAS6-AS1 gradually decreases in the samples from stage IV to stage I. **f** The expression score of GAS6-AS1 gradually decreases in the samples from T4 to stage T1. **g** Kaplan-Meier analysis shows that the high level of GAS6-AS1 correlates with poorer overall survival in all patients. **h** The patients with high level of GAS6-AS1 show a poor disease free survival in all patients. **i** ROC analysis confirms that GAS6-AS1 level is a sensitive indicator for predicting patient prognosis. **j** The schematic diagram of PDX model construction. **k** The clinicalpathological features of two patients for PDX construction. **l** Images of two PDX models. **m** When tumor volume reached 150 mm^3^, 5-FU was injected intraperitoneally every 3 days and ASO targeted GAS6-AS1 was injected intratumorally every 3 days. Tumor volumes from patient #8 were measured every 7 days. **n** Tumor volumes from patient #10 were shown. **o** Tumor weights for patient #8 and #10 were shown. **p** Representative images of IHC results of MCM3, KI67, Cyclin D1 and CDK4 in each group of patient #8 and #10. *P < 0.05; **P < 0.01; ***P < 0.001.

To test the practicality of GAS6-AS1 as combination treatment in CRC, ASO targeted GAS6-AS1 (Fig EV8) and PDX model (Fig 8j) were constructed. Clinical characterization of the donator patients is listed (Fig 8k). ASO of GAS6-AS1 combined with 5-FU decreased the volume and weight of tumor from two patients (Fig 8l-o). IHC results also indicated that ASO of GAS6-AS1 combined with 5-FU significantly decreased the expression of MCM3, KI67, cyclin D1 and CDK4, compared with that with only 5-FU (Fig 8p). Taken together, GAS6-AS1 might serve as a powerful biomarker and potential therapeutic target for combination drug therapy in CRC.

## Discussion

Chemotherapy resistance poses a formidable challenge to both adjuvant and neoadjuvant chemotherapy in advanced CRC. Emerging evidence implicates lncRNAs in the intricate mechanisms underlying resistance to various drugs. In this study, a comprehensive approach involving RNA-seq and WGCNA identified GAS6-AS1 as a closely associated lncRNA with TRG in CRC. The analysis of two distinct sample sets employing diverse methodologies consistently revealed a significant elevation in the expression of GAS6-AS1 in CRC. Strikingly, this upregulation of GAS6-AS1 exhibited a positive correlation with chemotherapy resistance, clinicopathological features, overall survival, and disease-free survival. GAS6-AS1 demonstrated potent oncogenic activity and played a pivotal role in 5-FU resistance in vitro and in vivo. Furthermore, GAS6-AS1 emerges as a sensitive indicator, holding promise for predicting prognosis in CRC. In conclusion, GAS6-AS1 is significantly upregulated, promotes 5-FU resistance and can act as an independent prognostic factor in CRC.

Previous studies have consistently reported the upregulation of GAS6-AS1 in various cancer types, showcasing its role in promoting cell growth, metastasis, and cell cycle transitions. In gastric cancer, GAS6-AS1 acts as an oncogenic lncRNA, contributing to increased cell growth and migration(Zhang, Dong et al., 2019). Similarly, in clear cell renal cell carcinoma, GAS6-AS1 is significantly elevated, leading to enhanced proliferation, metastasis, and glycolysis(Guo, Li et al., 2021). Moreover, in CRC, GAS6-AS1 has been identified as an upregulated lncRNA, promoting cell growth, invasion, and epithelial-mesenchymal transition(Chen, Zhou et al., 2022). While previous research has established GAS6-AS1’s involvement in various cancer-related processes, its specific role in CRC resistance and progression had not been previously elucidated. LncRNAs often exert their biological functions through interactions with regulatory proteins, miRNAs, or other cellular factors(Liu, Han et al., 2022, Liu, Shi et al., 2022, Zuo, Chen et al., 2020). GAS6-AS1 has been reported to act as a miRNA sponge in breast cancer, enhancing SOX9 expression(Wu, Xu et al., 2022). Additionally, it boosts MYC transactivation by directly binding Y-box binding protein 1 (YBX1) in acute myeloid leukemia, thereby promoting cell growth, colony formation, and cell cycle G1/S transition while reducing apoptosis(Zhou, Liu et al., 2021). In the present study, a series of RNA pulldown and RIP assays uncovered a direct interaction between GAS6-AS1 and PCBP1. As a member of hnRNPs, PCBP1 is involved in various aspects of nucleic acid metabolism, including transcriptional control, alternative splicing, mRNA stabilization, and translation regulation(Chaudhury, Chander et al., 2010, Grelet & Howe, 2019). PCBP1 has been reported to take part in various fields of malignant tumor progression in the role of tumor promotor or suppressor(Grelet, Link et al., 2017, Lv, Sun et al., 2022, Woosley, Dalton et al., 2019). As we discovered, PCBP1 rescued the regulations of GAS6-AS1 on 5-FU resistance, cell growth and cell cycle G1/S transition in CRC. Accordingly, the study not only expands our understanding of GAS6-AS1’s role in CRC but also unveils a novel molecular mechanism involving the GAS6-AS1/PCBP1 interaction. This discovery sheds light on potential therapeutic targets and offers insights into the intricate regulatory networks governing 5-FU resistance and malignant phenotypes in CRC.

In result of the subcellular distribution of both GAS6-AS1 and PCBP1 mainly on cytoplasm, GAS6-AS1 might work at the post transcriptional level. Subsequent RNA-seq and bioinformatic analyses revealed enrichment of biological functions related to the DNA replication and damage repair pathway. Within the genes associated with the DNA replication and damage repair pathway, the mRNA and protein levels of MCM3 were influenced by both GAS6-AS1 and MCM3. Further truncated RNA pulldown and RIP assays confirmed a genuine interaction and identified specific binding sites of PCBP1 with MCM3 mRNA. As a consequence, it was not surprising that GAS6-AS1 inevitably played a role in the binding of PCBP1 with MCM3 mRNA, which was called as ‘guider’. The following experiments showed that GAS6-AS1 enhanced the binding of PCBP1 with MCM3 mRNA. As we already know, the main function of 3’-UTR is to maintain the stability of mRNA. Our results also suggested that both GAS6-AS1 and PCBP1 extended the half-life of MCM3 mRNA. MCM3, a member of the minichromosome maintenance proteins (MCMs), is crucial for regulating DNA replication and maintaining normal cellular processes(Hatoyama & Kanemaki, 2023, Sedlackova, Rask et al., 2020, Wang, Chen et al., 2020, Xia, Sonneville et al., 2023). Dysregulation of MCMs often occurs in cancers, influencing tumor initiation, progression, and chemotherapy resistance by modulating cell cycle and DNA damage repair(Ha, Shin et al., 2004, Han, Mayca Pozo et al., 2015, Yang, Xie et al., 2019). Our results indicated that MCM3 rescued the regulation of GAS6-AS1 on 5-FU resistance and cell growth. In addition, GAS6-AS1 promoted MCM3 expression, while PCBP1 rescued its promotion on MCM3 expression. To summarize, GAS6-AS1 could serve as a ‘guider’ to enhance the binding of PCBP1 with the 3’-UTR of MCM3 mRNA, thereby increasing the stability and expression of MCM3 mRNA (Fig EV9).

Considering the significant regulation of GAS6-AS1 on 5-FU resistance in CRC, we next explore the possibility of combination therapy targeted on GAS6-AS1 and 5-FU in vivo. PDX model doesn’t undergo the screening process of in vitro culture, could maintain the original characteristics and heterogeneity of tumors, sustain molecular biology stability and preserve the original non tumor matrix and microenvironment of tumor tissue(Heinrich, Mostafa et al., 2021, Jin, Yoshimura et al., 2023, Zanella, Grassi et al., 2022). Thereby, PDX model was suitable for evaluating drug efficacy. In our PDX models, ASO targeted GAS6-AS1 combined with 5-FU got better efficacy in tumor inhibiting compared with 5-FU alone. Further clinical trials should be designed to test the clinical effects targeted GAS6-AS1. Also, the mechanisms underlying upregulation of GAS6-AS1 in CRC tissues and 5-FU resistant cells haven’t been explored. These questions may be pivotal in the process of developing more powerful drugs targeted GAS6-AS1. Whether GAS6-AS1 plays a similar role in the resistance of oxaliplatin and irinotecan remains unclear and needs further study, which contributes to augment the clinical applications of GAS6-AS1 target.

In summary, GAS6-AS1 is significantly upregulated, promotes 5-FU resistance and can act as an independent prognostic factor in CRC. Mechanistically, GAS6-AS1 enhances the stability and translation of MCM3 mRNAs by boosting its binding with PCBP1, thereby promoting cell growth and cell cycle G1/S progression. Moreover, PDX model confirmed the effects of targeting GAS6-AS1 on the 5-FU resistance in vivo. Our research revealed that GAS6-AS1 plays a crucial role in 5-FU resistance, and might serve as a powerful biomarker and potential therapeutic target for combination drug therapy in CRC.

## Materials and methods

### Data collection

Three CRC datasets from Fudan University Shanghai Cancer Center (FUSCC) constituted the basis of this study. A cohort of 22 advanced CRC patients, who underwent 5-FU-based neoadjuvant chemotherapy prior to surgery, was assembled. Postoperative pathology categorized 5 cases as TRG 0, 3 cases as TRG 1, and 14 cases as TRG 3. To further assess the expression of GAS6-AS1, 316 paired fresh frozen CRC tissues were meticulously selected and analyzed using reverse transcription polymerase chain reaction (RT-PCR). Additionally, a total of 295 paired paraffin-embedded CRC samples contributed to the construction of a tissue microarray (TMA), stored at −20 °C. The clinicopathological features and prognosis associated with these samples were systematically documented. The diagnoses of all clinicopathological features were independently confirmed by two pathologists. Notably, ethical approval for the present study was granted by the Ethics Committee of Fudan University Shanghai Cancer Center.

### Cell culture and plasmid construction

The HCT15 and HCT8 CRC cell lines, along with the 293T cell line, were procured from the Type Culture Collection of the Chinese Academy of Science (Shanghai, China). All cell lines were cultured in DMEM medium (Gibco, USA), supplemented with 10% fetal bovine serum (FBS; Hyclone, USA), and 1% penicillin-streptomycin, maintaining a 37 °C environment with 5% CO_2_. To establish 5-FU resistance, HCT15 and HCT8 cells were successfully cultured in the aforementioned medium supplemented with 5 µg/ml of 5-FU (Beijing Tsingke Biotech Co., Ltd., Beijing, China) and subsequently identified through IC50 detection.

For the regulation of GAS6-AS1, PCBP1, and MCM3 expression, oligonucleotides and overexpressed plasmids were meticulously constructed. The specific sequences of short hairpin RNAs (shRNAs) and antisense oligonucleotides (ASOs) can be found in Table EV5.

### RNA extraction and real-time quantitative polymerase chain reaction (RT‒ qPCR)

For the extraction of total RNA from cells and tissues, TRIzol (TaKaRa, Japan) was utilized. Subsequently, 500 ng of total RNA underwent reverse transcription to generate cDNA using the primeScript^TM^ RT Master Mix reagent kit (TaKaRa, Japan). Quantitative assessment of relative RNA expression was performed via RT-qPCR using SYBR Premix Ex Taq^TM^ (TaKaRa, Japan). The reference gene, β-actin, served as the internal control. The 2^-ΔΔCt^ method was employed for the analysis of relative expression levels, providing a normalized measure of gene expression. Specific primer sequences utilized for qPCR are detailed in Table EV5, facilitating precise and targeted amplification in the experimental process.

### Western blot

Western blot assays were carried out as previously described(Zhu, Yu et al., 2019). The following primary antibodies were used: PCBP1 (1:1000, Abcam, USA); MCM3 (1:1000, Abcam, USA); Cyclin D1 (1:1000, Abcam, USA); CDK4 (1:5000, Abcam, USA); GAPDH (1:5000; ABclonal, China). The following secondary antibodies were used: Anti-Rabbit (1:5000; ABclonal, China); Anti-Mouse (1:5000; ABclonal, China).

### CCK8, colony formation, edu and cell cycle assays

For CCK8 assays, 1000 cells/well were seeded in the 96-well plate. After 24 hours of incubation, 5 µg/ml 5-FU was introduced. Subsequently, 10 µl/well of CCK8 solution (NCM Biotech, China) was added. Following a continuous incubation period of 2 hours, absorbance was measured at 450nm. This process was repeated for three consecutive days to monitor absorbance changes. For colony formation assays, 600 cells/well were seeded in the 6-well plate. After a two-week incubation period, the culture medium was removed, and plates were washed with Phosphate Buffered Saline (PBS). Fixation was performed with 4% polymethanol for 15 minutes, followed by staining with 0.1% crystal violet for an additional 15 minutes. Finally, plates were photographed, and colony counts were calculated. The edu assay was performed by EdU Cell Proliferation Kit with Alexa Fluor 488 (Beyotime Biotechnology, China), following previously described procedures(Zhu, Guo et al., 2023). For cell cycle assays, cells were collected and fixed overnight in precooled 70% ethanol. After washing with 1 ml of precooled PBS, cells were stained with 10 μl of RNase A and 25 μl of propidium iodide (PI) staining solution in 500 μl of dye buffer at room temperature for 30 minutes. The cell cycle distribution was then determined using flow cytometry. All experiments were conducted in triplicate to ensure consistency and reliability of the results.

### Fluorescence in situ hybridization (FISH)

Cy3-labeled GAS6-AS1 and 18S probes, synthesized by RIBOBIO (Guangzhou, China), were employed for FISH assays in cells and TMA. The FISH assays were conducted using the FISH Kit (RIBOBIO, Guangzhou, China) following previously established protocols. GAS6-AS1 staining intensity was scored on a scale of 0 (negative staining), 1 (weak staining), 2 (moderate staining), and 3 (strong staining). The percentage of positively stained cells was scored as follows: 0 (no stained cells), 1 (1∼ 24%), 2(25 ∼ 49%), 3 (50 ∼ 74%), 4(75 ∼ 100%). The final score was calculated as the product of the intensity and percentage scores. Samples were categorized into two groups based on the final score: the low expression group (score 0–4) and the high expression group (score 6–12).

### RNA pulldown assays

RNA pulldown experiments were executed using the Pierce™ Magnetic RNA-Protein Pull-Down Kit (Thermo Scientific, Rockford, USA) according to the manufacturer’s instructions. Biotin-labeled RNAs for in vitro experiments were transcribed using the RiboTM RNAmax-T7 Biotin labeled Transcription Kit (RIBOBIO, Guangzhou, China). Briefly, cells were lysed, and the supernatant was collected after centrifugation. A portion (30 μl) of the supernatant was reserved as input, while the remaining lysate was subjected to RNA pulldown. After incubation with streptavidin magnetic beads, the beads were washed and boiled in SDS loading buffer for subsequent Western blot analysis.

### RNA immunoprecipitation (RIP) assay

RIP assays were performed using the Magna RIP^TM^ RNA-binding protein immunoprecipitation kit (Millipore, Massachusetts, USA). Cells in 15 cm dishes were lysed in complete RIP lysis buffer, and the lysate was stored at −80 °C. Antibodies (5 μg) were added to magnetic beads and incubated with rotation. Beads were then washed, and cell lysates were added, followed by overnight incubation at 4 °C. After washing, proteins bound to beads were digested with proteinase K buffer, and RNA was purified. RT-PCR was subsequently performed to assess the expression of specific RNA. Input samples were used as controls during the RIP assay.

### Luciferase reporter assay

Luciferase reporter plasmids, encompassing the 3’-UTR of MCM3 mRNA and a mutant sequence in the binding site with PCBP1, were synthesized by HarO Life Co. (Shanghai, China). These plasmids were co-transfected into cells along with overexpression or shRNA plasmids using Lipofectamine™ 2000 reagent. After 48 hours, firefly luciferase and renilla luciferase activity were measured using the Dual-Luciferase Reporter Assay System (Promega, Wisconsin, USA). The relative luciferase activity was calculated as the ratio of firefly luciferase to renilla luciferase activity. Each experiment was independently replicated in triplicate.

### RNA stability assay

To assess RNA stability, cells were treated with 5 μg/ml Actinomycin D (ACD) (Millipore, Massachusetts, USA) to inhibit de novo transcription. The level of MCM3 mRNA was measured by RT-PCR at four-hour intervals. Each experiment was conducted three times for robust results.

### Subcutaneous xenograft and PDX model

For the subcutaneous xenograft model, 5×10^6 CRC cells in 150 μL PBS were subcutaneously injected into the right flanks of male BALB/c nude mice (4 weeks old). Visible tumors, 10 days post-injection, were treated with 50 mg/kg 5-FU intraperitoneally, twice weekly. Tumor volume was measured weekly. After 4 weeks of 5-FU treatment, mice were sacrificed, and tumors were subjected to weight measurement and histological analysis.

The patient-derived tumor xenograft (PDX) model utilized 4-week-old male NOD/SCID (NCG) mice. CRC tissues, obtained post-surgical resection, were subcutaneously implanted into the right flanks of NOD/SCID mice. When tumors reached a volume of 150 mm^3, 50 mg/kg 5-FU was intraperitoneally administered and ASO was intratumorally injected, twice weekly. Tumor volume was monitored weekly, and after 5 weeks of 5-FU and ASO treatment, mice were sacrificed for weight measurement and histological analysis. All in vivo experiments received approval from the Animal Care Committee of Fudan University Shanghai Cancer Center (FUSCC-IACUC-2023038).

### Statistical analyses

Statistical analyses were performed using GraphPad Prism 9.0 and R software. Student’s t-test was used for comparing two groups, while one-way or two-way ANOVA was applied for multi-group comparisons. The χ2 test and Fisher’s exact test were employed for counting data differences. Survival analysis utilized the Kaplan-Meier method, and significance was evaluated using the log-rank test. A p-value < 0.05 was considered statistically significant.

## Data availability

The online datasets are available in The Cancer Genome atlas (https://portal.gdc.cancer.gov/) and Gene Expression Omnibus (https://www.ncbi.nlm.nih.gov/geo/) databases. The other data used for the study is available from the corresponding author on reasonable request.

## Acknowledgements

This study was supported by the Science and Technology Commission of Shanghai Municipality (20DZ1100101), Shanghai Hospital Development Center (SKXZ2028), and National Natural Science Foundation of China (82003060). The research was approved by the Ethics Committee of Fudan University Shanghai Cancer Center (2112248-23). All in vivo experiments were approved by the Animal Care Committee of Fudan University Shanghai Cancer Center (FUSCC-IACUC-2023038).

## Author contributions

Zhonglin Zhu: Investigation; Methodology; Writing – original draft. Ye Xu: Resources; Conceptualization; Project administration; Funding acquisition; Supervision; Data curation. Minghan Li: Methodology; Writing-review and editing; Investigation. Junyong Weng: Investigation; Funding acquisition. Shanbao Li: Data curation; Methodology. Tianan Guo: Investigation; Methodology. Yang Guo: Methodology; writing-review and editing. All of the authors discussed and approved the final manuscript.

